# SINGLE CELL RESOLUTION OF THE ADULT ZEBRAFISH INTESTINE UNDER CONVENTIONAL CONDITIONS, AND IN RESPONSE TO AN ACUTE *VIBRIO CHOLERAE* INFECTION

**DOI:** 10.1101/2023.04.14.536919

**Authors:** Lena O. Jones, Reegan J. Willms, Mckenna Eklund, Ralph Derrick V. Graham, Xinyue Xu, Minjeong Shin, Edan Foley

## Abstract

*Vibrio cholerae* is an aquatic bacterium that primarily infects the gastrointestinal tract, causing the severe and potentially deadly diarrheal disease, cholera. Despite the impact of *Vibrio* on global health, our understanding of host mucosal responses to the pathogen at the site of infection remains limited, highlighting a critical knowledge gap that must be addressed to develop more effective prevention and treatment strategies. Using a natural infection model, we combined physiological and single-cell transcriptomic studies to characterize adult zebrafish guts raised under conventional conditions and after a challenge with *Vibrio*. We discovered that *Vibrio* causes a mild mucosal immune response characterized by T cell activation and enhanced antigen capture in the epithelium. Additionally, we discovered that *Vibrio* suppresses host interferon signaling, and that ectopic activation of interferon significantly alters the course of infection. Notably, we also found that the adult zebrafish gut shares many similarities with mammalian counterparts, including the presence of previously undescribed Best4+ cells, tuft cells, and a population of basal cycling cells. These discoveries provide important insights into host-pathogen interactions and emphasize the utility of zebrafish as a natural model of *Vibrio* infection.

## INTRODUCTION

The aquatic bacterium *Vibrio cholerae* (*Vc*) is a non-invasive, deadly human pathogen that infects millions, claiming roughly 100,000 lives each year through the diarrheal disease cholera, with disproportionately high mortality rates among children^1^. Although rehydration can prevent death, access to clean drinking water is frequently limited during cholera outbreaks, especially in regions impacted by civil conflict or natural disaster^2,3^. Antibiotics and vaccines are effective against *Vc*, but antibiotic-resistant strains have emerged^4^, and the efficacy of existing vaccines wanes with time^5,6^. Despite the importance of developing therapies that protect from disease, we have limited understanding of the immediate impacts of *Vc* on immune responses within the host gut. A paucity of information on intestinal anti-*Vc* responses significantly impedes our ability to develop rational, effective therapies or markers for disease severity. Gene expression data and histological examination of patient biopsies indicate a mild mucosal immune response that likely includes granulocyte engagement and T cell mobilization^7–9^. However, assigning host responses directly to *Vc* is complicated. Many patients were co-infected with intestinal parasites, treated with antibiotics, and may have had prior *Vc* exposures. As a result, the intestinal response to a natural, primary encounter remains almost entirely unknown.

Several experimental models have been deployed to characterize the impacts of *Vc* on a host gut^10^. *Drosophila melanogaster* is excellent for studying *Vc* pathogenesis and pathogen-commensal interactions during disease^11–14^. However, flies lack immune-regulatory Intestinal Epithelial Cell (IEC) types found in humans, and the fly gut does not have associated populations of lymphoid and myeloid cells. Rabbits and mice are valuable mammalian models^15–17^, but require work with immunologically immature juveniles, exposure to antibiotic cocktails, or surgical manipulation to replicate the diarrheal disease observed in humans^18^.

In recent years, the zebrafish, *Danio rerio*, has emerged as an excellent vertebrate model of *Vc* pathogenesis^19–21^. Fish offer several advantages that other infection models lack. Wild and lab-raised fish naturally co-associate with *Vibrio* strains, and fish are a candidate vector for the transmission of pathogenic *Vc* in the wild^22,23^. Of equal significance, the simple addition of live *Vc* to fish water results in a host disease hallmarked by rapid diarrheal shedding of live pathogen^24^. Thus, the fish is ideal for studies that characterize host responses to *Vc* in a natural infection. Despite general progress with the fish-*Vibrio* model, we have a rudimentary understanding of the gut immune response to infection. Physiologically, the fish intestine appears quite similar to mammalian counterparts^25^. However, critical gaps in our appreciation of adult intestinal physiology impair our ability to understand how *Vc* impacts gut defenses. Developmental, transcriptional and functional studies have outlined broad similarities between fish and mammalian intestines^26–33^. However, we have an incomplete picture of adult fish IEC composition and arrangement within the gut, and we lack a detailed characterization of the identity and state of gut-associated leukocytes. We believe a comprehensive characterization of cell functions encoded within the adult gut is essential for developing the zebrafish as a *Vibrio* infection model.

To advance our understanding of *Vc* pathogenesis in a natural infection, we completed structural and transcriptional characterizations of the adult fish gut in the presence or absence of pathogenic *Vc*. We discovered previously unknown IEC types, including lineages with human counterparts such as tuft cells, and Best4+ absorptive cells. We simultaneously generated an unbiased atlas of the transcriptional states of gut-associated leukocytes that included granulocytes, macrophages, dendritic cells, B cells, and specialist T cell subpopulations. We resolved the transcriptional response of each cell type to *Vc* at the level of individual cells and discovered that *Vc* prompts a moderate immune response in mature epithelial cells, activates protective anti-bacterial responses in gut-associated leukocytes, and suppresses expression of key interferon pathway elements throughout the gut. We consider possible relationships between infection and interferon signaling noteworthy, as interferon is the subject of natural selection in a human population from a cholera-endemic region^34^. In follow-up work, we showed that pre-treatment of infected adults with an interferon agonist reprogrammed the transcriptional response to infection and significantly modified host colonization, suppressed epithelial death, and increased diarrheal shedding of *Vc* by the host, suggesting a link between interferon and intestinal disease. Our work expands the value of zebrafish for studying enteric pathogens and resolves the gut response to *Vc* in a natural infection model. We believe a detailed resolution of the intestinal immune response to infection will be particularly valuable for the development of effective vaccines and biomarkers for disease.

## RESULTS

### The Adult Intestine Associates with Sub- and Intraepithelial Leukocytes

To determine how *Vc* affects intestinal function and morphology in adult fish, we first used tissue histology, immunofluorescence, and immunohistochemistry to characterize gut architecture in wildtype adults. H&E stains confirmed earlier reports that the adult intestine includes an epithelial monolayer that protrudes into the lumen^25^ (Figure 1A). Each fold contained physically distinct cell types that included mucus-filled goblet cells, columnar epithelial cells, and inter-epithelial leukocytes (Figure 1B-C). Apical intestinal epithelial cells (IECs) were marked by membrane-enriched beta-catenin staining, while basal goblet cells displayed a nuclear enrichment of beta-catenin signal (Figure 1D-E), possibly reflecting a role for beta-catenin in secretory cell specification as described previously for mammals^35–37^. We also detected basal clusters of PCNA-positive proliferating cells that parallel a recent description of cycling cells in the adult gut and are highly reminiscent of the intestinal stem cell/transit amplifying cell compartment of mammalian small intestinal crypts (Figure 1F). Further emphasizing similarities between zebrafish and mammals, we also detected an apical accumulation of pSMAD-positive cells (Figure 1G), indicating a basal to apical gradient of Bone Morphogenetic Protein activity in IECs.

**FIGURE 1:**
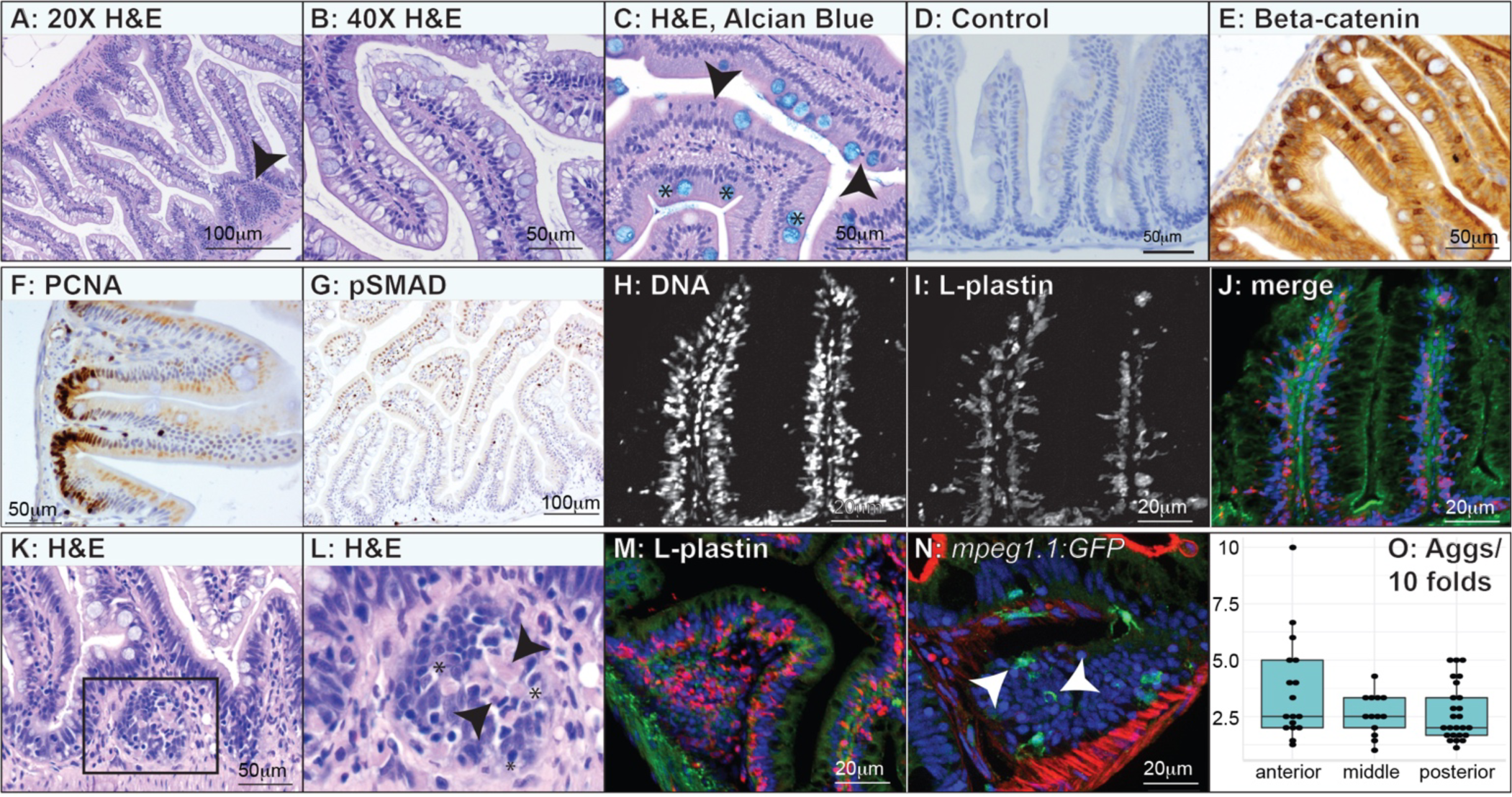
Arrangement of Epithelial Cells and Associated-Leukocytes in the Adult Zebrafish Gut. (A-C) H&E stains of sagittal posterior intestinal sections from adult zebrafish. Scale bars are indicated in each panels. In A, a prominent subepithelial leukocyte aggregate is marked with an arrowhead. In C, tissue is also stained with alcian blue to visualize goblet cells. Representative inter-epithelial leukocytes and alcian blue-positive goblet cells are marked with arrowheads and asterisks, respectively. (D-G) Immunohistochemical images of a sagittal posterior intestine section stained without a primary antibody (D, control), for beta-catenin (E), PCNA (F), and pSMAD (G). (H-J) Immunofluorescence imaging of two posterior intestinal folds stained for DNA (H) and the pan-leukocyte marker, L-plastin (I). In the merged, false-colored image (J), DNA is labeled blue, leukocytes are labeled red, and the tight junction marker ZO-1 is labeled green. (K) H&E stain of a subepithelial leukocyte aggregate from the posterior intestine. (L) Digital magnification of the boxed region from K. Round, lymphoid-like cells are marked with asterisks and larger, eosin-rich cells are marked with arrowheads. (M) A merged, false-colored image of a sub-epithelial aggregate with DNA labeled blue, leukocytes labeled red, and the tight junction marker ZO-1 labeled green. (N) Immunofluorescence imaging of a leukocyte aggregate from *mpeg1.1:GFP* fish with DNA labeled blue, and filamentous actin labeled red. GFP-positive APCs within a lymphoid aggregate are labeled with a white arrowhead. (O) Quantification of sub-epithelial lymphoid aggregates per ten folds in serial sections prepared from five fish.

Immunofluorescence staining of the pan-leukocyte marker L-plastin revealed a regular distribution of intraepithelial leukocytes (Figure 1H-J), showing that, like mammals, fish IECs interdigit with specialist immune cells. We also observed occasional subepithelial aggregates of intestinal leukocytes (Figure 1A, K-M) that included round, lymphoid-like cells, large, eosin-rich cells (Figure 1L), and cells that expressed the antigen-presenting cell marker *mpeg1.1* (Figure 1N). On average, we detected one leukocyte aggregate per four folds throughout the adult intestine (Figure 1O). In combination, our data highlight the adult fish gut as an elaborate structure with overt similarities to mammalian counterparts, including subepithelial aggregates of gut-associated leukocytes.

### The Intestine Contains a Complex Population of Spatially and Functionally Specialist Epithelial Cells

To uncover the range of cellular functions encoded within such a complex epithelial structure, we resolved the transcriptomes of single cells purified from healthy adult fish guts. We successfully generated high-quality expression data for 18,358 cells that partitioned into 16 transcriptional states through graph-based clustering (Figure 2A). We assigned a cellular identity to each state based on expression of established cell type markers from fish and mammals (Figure 2B, Supplementary table 1). Among the transcriptional clusters, we identified leukocytes, differentiated stromal cells (marked by *col1a1b*, *vim*, *kdrl*), and nine distinct IEC types. Two IEC types that we have provisionally labeled progenitor 1 and 2 (Figure 2A-B), were characterized by prominent expression of genes associated with proliferation (e.g. *mki67*, *pcna*, *ccnd1*, Figure 2C), alongside markers of Notch pathway activity (e.g. *dld*, *ascl1a*), including *her15.1* (Figure 2D), a pan-stem cell marker in zebrafish^38–40^. The progenitor two population also expressed the recently identified ISC marker, *prmt1* (Figure 2E), adding support to the hypothesis that *prmt1* marks adult stem cells in zebrafish. Fluorescence in situ hybridization confirmed that *her15.1*+ cells are proliferative residents of the interfold base (Figure 2F), with 55% of all *her15.1*-positive cells incorporating the S-phase label EdU (Figure 2F). We believe our transcriptional identification of cycling cells with enriched Notch pathway activity that express stem cell markers (Figure 2A-E), alongside our immunohistochemical characterization of multiple proliferative cells at each fold base (Figure 1E) suggests that, like mammals, the adult fish intestine contains two populations of basal, cycling, progenitor cells. However, we emphasize that additional work is needed to unambiguously determine if the cycling populations directly correspond to ISC/transit amplifying cells observed in mammalian intestinal crypts.

**FIGURE 2:**
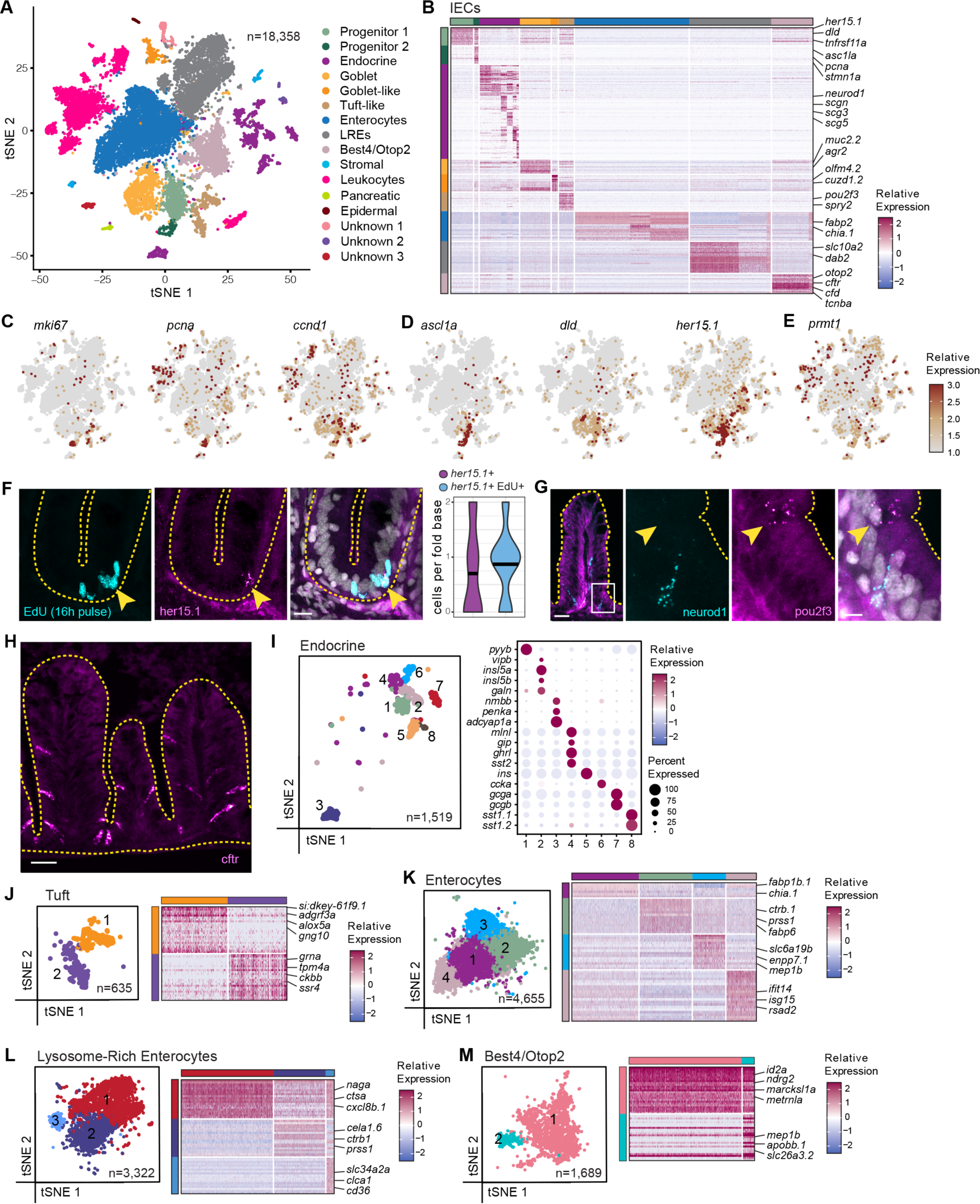
The Intestinal Epithelium Contains a Complex Mix of Proliferative and Mature Specialist Epithelial Cells. (A) 2D t-SNE projections of 18,358 intestinal cells color coded by cell type. (B) Heatmap of IEC cluster markers colored by relative gene expression and arranged according to cell type. Cell types are indicated by colored bars on the left and top. Prominent markers for each cluster are shown on the right y-axis of the heatmap. (C-E) Feature plots showing relative expression of proliferation markers (C), Notch pathway components (D) and the stem cell marker *prmt1* (E). (F) Fluorescence visualization of EdU-positive cycling cells (cyan), and *her15.1* positive cells (magenta) in an intestinal fold base. In the merged image all nuclei are labeled in white and a *her15.1*/EdU double-positive cell is marked with a yellow arrowhead. Quantification shows the number of *her15.1* single-positive and *her15.1*/EdU double positive cells per fold base. Scale bar = 10μm (G) Fluorescence visualization of *neurod1*-positive endocrine cells (cyan), and *pou2f3*-positive tuft cells (arrowhead, magenta) in an intestinal fold. In the merged image all nuclei are labeled in white. Scale bar = 25μm for left panel and 5μm for inset. (H) Fluorescence *in situ* hybridization showing the basally enriched expression pattern of the Best4+ cell marker *cftr* in three intestinal folds. Scale bar = 25μm. (I) t-SNE plot of enteroendocrine cells color coded by subset type. Bubble plot shows the relative expression levels for eighteen peptide hormones across all eight cell types (J-M) t-SNE plots of Tuft (J), Enterocyte (K), Lysosome-Rich Enterocytes (L) and Best4+ clusters (M) from original graph-based analysis, re-clustered and color coded by cell type. In each instance, heatmaps show expression of subset cell markers colored by relative gene expression. Cell types are indicated by colored bars on the left and top. Markers for each subset are shown on the right y-axis of each heatmap.

We also identified transcriptional profiles for secretory enteroendocrine (*neurod1+*, *scgn+*, *nkx2.2a+*) goblet (*agr2+*, *CABZ0180550.1+*) and tuft cells (*pou2f3+*, *spry2+*, *alox5a+*), as well as absorptive enterocytes (*fabp2+*, *cd36+*, *chia.1+*), lysosome-rich enterocytes (LREs, *fabp6+*, *tmigd1+*, *slc10a2+*), and recently discovered Best4+ cells (*best4+*, *otop2+*, *cftr+*) (Figure 2B). We found that *pou2f3*+ IECs did not express the endocrine marker *neurod1* and were morphologically distinct from goblet cells (Figure 2G), confirming the existence of tuft cells in the adult fish. We then compared our adult transcriptional data with a single cell gene expression atlas we recently prepared for six-day post-fertilization larvae. In general, we observed broad overlaps between adult and larval data sets (supplementary figure 1A), suggesting functionally conserved cell types in the respective intestines. However, adults had a clear expansion of leukocytes populations (supplementary figure 1B), and a possible increase in goblet numbers (supplementary figure 1B), likely reflecting the emergence of lymphocyte-driven adaptive defenses in the adult.

To fully characterize IEC cell types, we systematically analyzed gene expression profiles of all secretory and absorptive lineage for specialist subsets. In this manner, we resolved the enteroendocrine population (Figure 2B) into eight subtypes based on their unique peptide-hormone expression profiles (Figure 2I). We also identified two tuft cell subsets distinguished by expression of *si:dkey-61f9.1*, a C-type lectin domain-containing protein that exhibits homology to the IgE receptor *fcer2* (Figure 2J). Upon examination of enterocytes, we identified four unique clusters. Cells within cluster one expressed markers of lipid metabolism, while cluter two cells were enriched for expression of the bile acid binding protein gene *fabp6*; cluster three cells expressed high levels of endopeptidases including meprin subunits; and cluster four cells expressed interferon-response genes, such as *ifit14*, *isg15* and *rsad2* (Figure 2K). Our identification of interferon-sensitive enterocytes in adults matches our recent discovery of an enterocyte population in conventionally raised larvae that expressed interferon response genes^41^, and suggests enhanced interferon signaling within a dedicated enterocyte population (Figure 2K). We also discovered transcriptionally distinct LRE and Best4+ populations (Figure 2L-M), where LRE cluster three and Best4+ cluster two closely resembled canonical enterocytes, suggesting that these absorptive subtypes may arise from a common progenitor. *In vivo* visualization of the Best4+ cell marker *cftr* showed that *cftr*-positive cells generally resided in the lower half of intestinal folds (Figure 2H), indicating a spatially restricted requirement for Best4+ cells. In sum, our imaging and transcriptional data reveal the adult fish gut as an integrated community of spatially organized specialist cells that act in concert to support animal health and protect from infection. We consider our findings of value to a large community of researchers and have made all conventional IEC and leukocyte gene expression data available as a searchable tool on the Broad Institute Single Cell Portal (https://singlecell.broadinstitute.org/single_cell/study/SCP2141/adult-zebrafish-intestine#study-visualize).

### Gut-Associated Leukocytes Mediate Critical Protective Responses

Given the importance of gut-associated leukocytes for intestinal homeostasis, and for defense against pathogenic microbes, we elected to characterize the transcriptional states of intestinal leukocytes in greater detail. Using graph-based clustering of 2985 leukocytes, we identified five cell types: a dominant population that we provisionally labeled T cells, alongside B cells, macrophages, granulocytes and dendritic cells (Figure 3A). Each cluster had a unique gene expression profile (Figure 3B, Supplementary Figure 2A-E, supplementary table 2), that indicate specialist, cell type-specific leukocytes contributions to intestinal innate and adaptive defenses (Supplementary Figure 2F).

**Figure 3:**
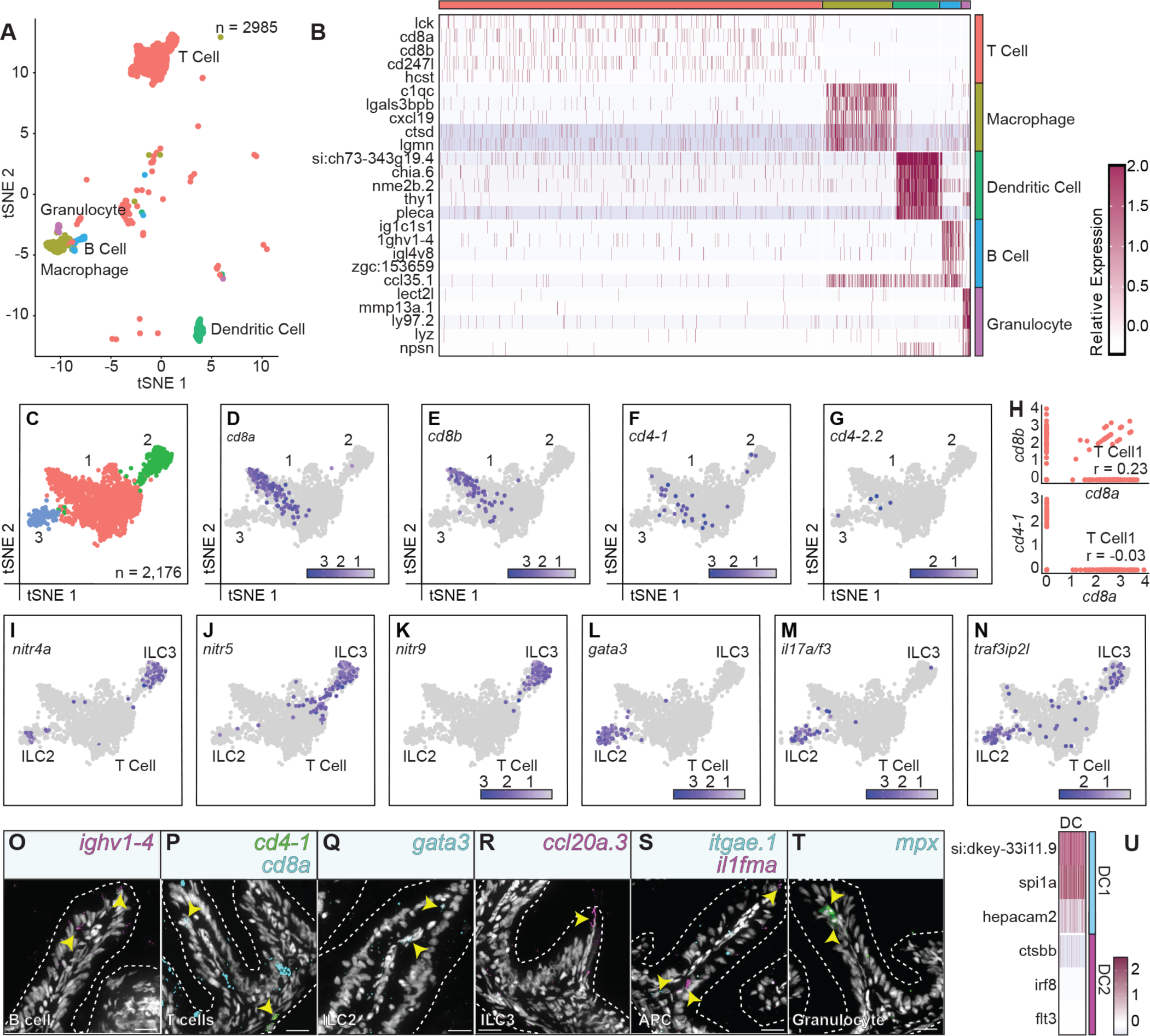
The Adult Intestinal Epithelium Associates with Protective Lymphoid and Myeloid Cells. (A) 2D t-SNE projections of 2,985 gut-associated leukocytes color coded by cell type. (B) Heatmap of leukocyte cluster markers colored by relative gene expression and arranged according to cell type. Cell types are indicated by colored bars on the right and top. Markers for each cluster are labeled on the y-axis of the heatmap. (C) t-SNE plots of T-like lymphoid cells color coded by cell subtype. (D-G) Feature plots showing the expression of *cd8a*, *cd8b*, *cd4-1* and *cd4-2.1*. (H) Scatter plots examining the relative expression of *cd8a* and *cd8b*, and *cd8a* and *cd4-1* across conventional T cells. r value indicates Pearson correlation for both curves. (I-M) Feature plots showing the expression of ILC3 marker genes *nitr4a*, *nitr5*, and *nitr9*, as well as ILC2 marker genes *gata3* and *il17a/f3*. Panel (N) shows the expression of a pan-ILC marker *traf3ip2l*. (O-T) Fluorescence *in situ* hybridization images showing intestinal distribution of B cells (*ighv1-4*+), T cells (*cd4-1* and *cd8a*+), ILC2s (*gata3*+), ILC3s (*ccl20a.3*+), macrophages (*itgae.1*+), dendritic cells (*il1fma*+) and granulocytes (*mpx*+). In each panel, the respective cell types are indicated with a yellow arrowhead. (U) Heatmap of the relative expression of conventional dendritic cell markers (DC1) and plasmacytoid dendritic cell markers (DC2). Conventional dendritic cell markers are specifically enriched in intestinal dendritic cells.

Looking at T cell gene expression profiles, we noted relatively modest enrichment for classical T cell markers such as *lck*, *cd8*, or the *CD3zeta-like* gene *cd247l*, suggesting heterogeneity within the T cell population. Upon re-clustering the T cell group, we resolved three unique transcriptional states (Figure 3C). The most prominent cell type was conventional T cells that expressed cd4 or cd8 paralogs (Figure 3D-G), indicating an abundance of cytolytic and helper T cells in adult fish guts. Intestinal T cells appeared to be fully mature, as we detected co-expression of *cd4* or *cd8* paralogs within individual cells (Figure 3H), but never co-expression of *cd4* and *cd8* (Figure 3H). The remaining two subsets were marked by expression of numerous *nitr* genes (Figure 3I-K) or *gata3* and *il17a/f3* (Figure 3L-M), respectively, matching the recently characterized type three and two populations of gut-associated innate lymphoid cells^42^ (ILCs, Figure 3P). Functionally, ILC2s and ILC3s displayed clear distinctions, with ILC3s primarily expressing genes linked with T cell activation and type 2 antibacterial responses, whereas ILC2s expressed genes associated with adaptive immune responses (Supplementary Figure 2G).

To chart the intestinal distribution of gut-associated leukocytes, we used cell type-specific *in situ* hybridization probes to visualize each cell type in the adult gut (Figure 3 O-T). Although more extensive mapping is needed, our data suggest a degree of spatial coordination among gut-associated leukocytes. For example, whereas B cells (*ighv1-4*+) appear to accumulate in apical folds (Figure 3O), *cd8*+ T cells have a relatively broad epithelial distribution (Figure 3P), and *cd4*+ T cells appear to accumulate basally (Figure 3P). Like other vertebrates, ILC2s (*gata3*+) and ILC3s (*ccl20a.3*+) cells associated with intestinal epithelial cells (Figure 3Q-R), while antigen presenting cells (*itgae.1*+ macrophages and *il1fma*+ dendritic cells) and granulocytes (*mpx*+) were more broadly distributed, including among non-epithelial cells (Figure 3R-T). A recent study demonstrated that zebrafish produce conventional antigen-processing dendritic cells, as well as plasmacytoid dendritic cells typically associated with anti-viral responses^43^. We found that gut-associated dendritic cells primarily expressed conventional markers such as the CD225 Dispanin family member *si:dkey-33i11.9*, *spi1a*, and *hepacam2*, with minimal expression of the plasmacytoid DC markers *ctsbb*, *irf8*, and *ftl3* (Figure 3U), indicating that intestinal dendritic cells primarily belong to the conventional subset. In combination, our transcriptional and spatial data point to cell type-specific execution of intestinal defenses by a mix of lymphoid and myeloid lineages.

### *Vibrio cholerae* Infections Modify Innate Immune Responses in the Adult Intestinal Epithelium

Thus far, our data reveal the structural and transcriptional organization of adult intestinal cells during homeostasis. To understand how infection with *Vc* impacts gut function, we generated single cell gene expression profiles of intestines from adult fish that we challenged for sixteen hours with *Vc* at the same time as our profiles of conventional, uninfected fish. For these experiments, we worked with the clinical V52 *Vc* isolate that colonizes and induces disease in fish^44,44,45^. We found that *Vc* associated with anterior, middle and posterior intestinal segments to approximately equal levels throughout three days of infection (Figure 4A-C). We did not observe visible signs of disease or distress in infected animals. However, histological stains revealed extensive epithelial damage during infection. Unchallenged fish had large, regular villus-like projections that permeated the intestinal lumen (Figure 4D-E). In contrast, infections disrupted epithelial integrity (Figure 4F), leading to breaches, lumenal shedding of intestinal material (Figure 4G), and occasional internalization of microbial cells with Vc-like morphology (Figure 4H). TUNEL stains revealed an increase in IEC death for infected fish, including prominent shedding of dying cellular material into the lumen (Figure 4H-I). Looking at differential gene expression within an integrated dataset derived from IECs of infected and unchallenged fish (Figure 4K), we observed immune responses to infection that included increased expression of proinflammatory genes (e.g. *grn1*, *agr2*, *icn* Figure 4L) and suppressed transcription of numerous interferon response signatures (e.g. *isg15*, *ifit15*, *rsad2*, Figure 4L-M). Examination of cell type specific responses (supplementary table 3) revealed modest suppression of genes required for leukocyte recruitment, most notably in Best4+ cells (Figure 4N); enhanced expression of genes involved in antigen presentation, particularly in enterocytes (Figure 4O); and cell type-specific shifts in expression of genes linked with mucin production, including an unexpected attenuation of *mucin2* expression in goblet cells (Figure 4P). The most prominent epithelial response to infection was a broad suppression of genes associated with the interferon response in mature, differentiated IECs. With the exception of progenitors, we found that exposure to *Vc* significantly attenuated expression of a host of interferon signaling markers (e.g. *ifi46*, *isg15*, *stat1a*, *stat1b* and *stat2*) in secretory and absorptive cells (Figure 4K). Thus, our data show that infection with pathogenic *Vc* impacts intestinal integrity, and prompts an epithelial response characterized by induction of genes required for antigen capture, alongside suppression of genes linked with interferon pathway activity.

**Figure 4:**
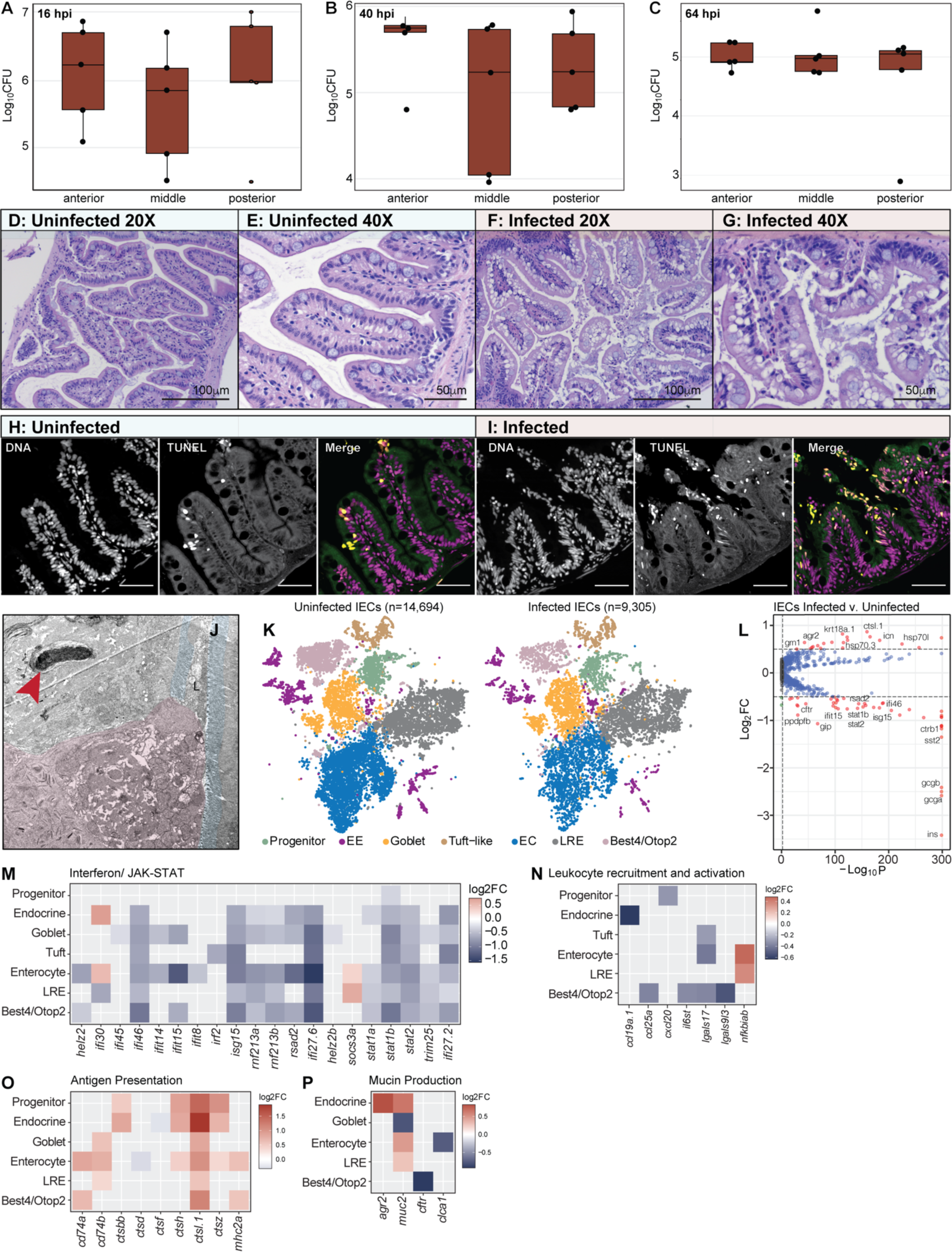
*Vibrio cholerae* activates Inflammatory Responses While Suppressing Interferon Signaling in IECs. **(A-C).** CFU Counts of gut-associated *Vc* in anterior, middle, and posterior intestinal sections of infected fish, 16h (A), 40h (B) and 64h post-infection (h.p.i.). (D-G) H&E stains of sagittal posterior intestinal sections from uninfected (D-E) and infected adult zebrafish (F-G) at the indicated magnifications. Scale bars are indicated in all panels. (H-I) Visualization of TUNEL-positive cells in the intestinal epithelium of uninfected fish (H), or fish challenged with *Vc* (I). In the merged images, TUNEL+ cells are yellow, and nuclei are magenta. (J) Transmission Electron Microscopy image from a sectioned, infected adult intestine. An internalized microbe with a *Vc*-like morphology is indicated with a red arrowhead. (K) t-SNE projections of profiled cells from an integrated data set generated from uninfected and infected IECs color-coded by cell type. The left panel shows uninfected IECs and the right panel shows infected IECs. (L) Volcano plot of differentially expressed genes in infected epithelial relative to uninfected controls. Y-axis shows the relative expression changes on a log2 scale, and x-axis shows significance values as a -log10 value. (M-P) Heatmaps showing differentially expressed genes related to interferon signaling (M), leukocyte recruitment (N), antigen presentation (O) and mucin production (P) in infected cells relative to uninfected counterparts.

### Characterization of the *Vibrio* Response in Gut-Associated Leukocytes

*Vc* is a non-invasive microbe that exclusively infects the intestine. However, we have minimal data on the impacts of *Vc* on leukocyte function at the site of infection in any experimental model, or in patient samples. For example, we do not know how intestinal antigen-presenting cells or lymphocytes respond to a primary encounter with *Vc*. Our integrated single cell data included 3,871 gut-associated myeloid and lymphoid cells (Figure 5E), providing a unique window into the leukocyte response to *Vc*.

**Figure 5:**
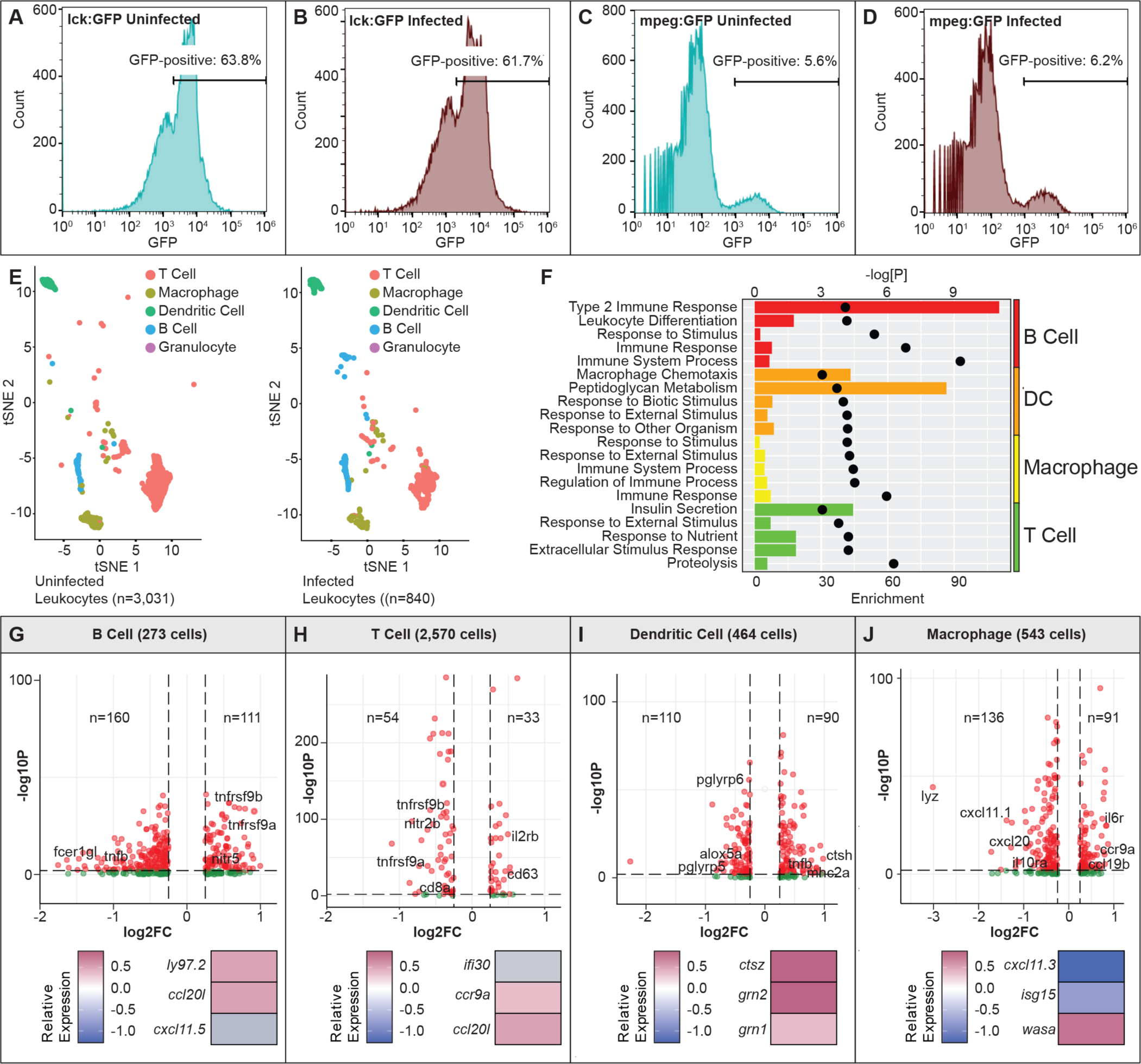
Lymphoid and Myeloid Responses to a Primary Encounter with *Vc*. (A-D) Percentage of gut-associated leukocytes that express GFP reporters under control of the *lck* or *mpeg1.1* promoters in uninfected fish and in fish twenty hours after infection. (E) t-SNE projections of an integrated data set generated from all leukocytes and color-coded by cell type. The left panel shows uninfected leukocytes and the right panel shows infected leukocytes. (F) The top five non-redundant Gene Ontology terms induced by infection per cell type. Enrichment score is represented by bar length and the negative log p value is shown with black circles. (G-J) Volcano plots of infection-dependent differentially expressed genes in the indicated cell type, with significance values represented on the y-axis and relative fold-change of gene expression on the x-axis. For all four cell types, only genes with log2 expression changes greater than 0.25 are shown. Heatmaps indicate expression values of three *Vc*-response genes for each cell type.

Infections did not have appreciable effects on the proportion of gut-associated leukocytes that expressed GFP reporters driven by the T cell-specific *lck* promoter (*lck:GFP*, Figure 5A-B), or the antigen-presenting cell-specific *mpeg1.1* promoter (*mpeg1.1:GFP*, Figure 5C-D). In contrast, we saw substantial, cell type-specific effects of *Vc* on gene expression in each leukocyte cell type. In general, the B cell response appeared more pronounced that other cell types (Figure 5G-J), possibly reflecting the dual roles of zebrafish B cells in antigen presentation and antibody production. For example, infection enhanced expression of antibacterial type two immune response genes in B cells (Figure 5F), as well as genes linked with T cell recruitment and activation, such as the *ccl20l* cytokine and the co-stimulatory paralogs *tnfrsf9a* and *tnfrsf9b* (Figure 5G). At this early stage of infection, the T cell response was muted relative to all other cell types (Figure 5H, K), likely reflecting the fact that T cells are dispensable for antigen capture during initial stages of infection. Many of the *Vc*-response genes in T cells were primarily linked with metabolic adaptations (Figure 5F). However, we observed signatures of T cell activation that included enhanced expression of the il2 receptor beta ortholog *il2rb*, and elevated expression of *ccr9a*, a gene linked with maturation of mucosal T cells (Figure 5H).

Looking at phagocytes, we detected lineage-specific responses to infection. For example, dendritic cells modified expression of genes required for peptidoglycan metabolism and cellular chemotaxis (Figure 5F), enhanced expression of the pro-inflammatory granulins *grn1* and *grn2* (Figure 5I), and increased expression of key components of the antigen processing and presentation machinery (e.g. *ctsh*, *ctsz*, *mhc2a*, Figure 5I). In agreement with a previous report, unchallenged macrophages expressed pro- and anti-inflammatory M1 and M2 markers at the population level^46^ (supplementary figure 3A-D). However, individual macrophages showed clear signs of polarization towards either the M1 or M2 phenotype (supplementary figure 3E-G), with a pro-inflammatory M1 profile dominating the response to infection (supplementary figure 3H-I). Upon infection, we observed altered macrophage-specific expression of immune regulators (Figure 5F), diminished expression of the anti-inflammatory il10 receptor, *il10ra*, and increased expression of *ccl19a*, a chemokine-encoding gene associated with T cell recruitment (Figure 5J).

In sum, our data uncover a coordinated engagement of specialist immune cells by *Vc* that promotes inflammatory defenses within the gut, increases the capacity to capture microbial antigens, and enhances recruitment and activation of gut-associated T cells. Notably, we discovered that, like the IEC response to infection, *Vc* attenuated expression of multiple interferon-response genes, including *cxcl11* paralogs in B cells and macrophages (Figure 5G, J), *ifi30* in T cells (Figure 5H) and *isg15* in macrophages (Figure 5J), indicating that suppressed interferon signaling is a unifying feature of the host intestinal response to *Vc*.

### *Vibrio cholerae* Suppresses Interferon Signaling in Infected Intestines

Our single cell gene expression data indicated an intestinal immune response to *Vc* characterized by pro-inflammatory gene expression, and attenuated interferon signals in IECs (Figure 4) and gut-associated leukocytes (Figure 5). Consistent with links between infection and diminished interferon activity, we detected significantly suppressed expression of the interferon-sensitive *viperin* ortholog *rsad2* in mature IECs, including endocrine, goblet, and enterocyte cells (Figure 6A). We also discovered that challenges with *Vc* greatly diminished the extent of STAT phosphorylation in IECs (Figure 6D-E) relative to unchallenged IECs (Figure 6B-C), further supporting an inhibitory effect of *Vc* on intestinal IFN signaling.

**Figure 6:**
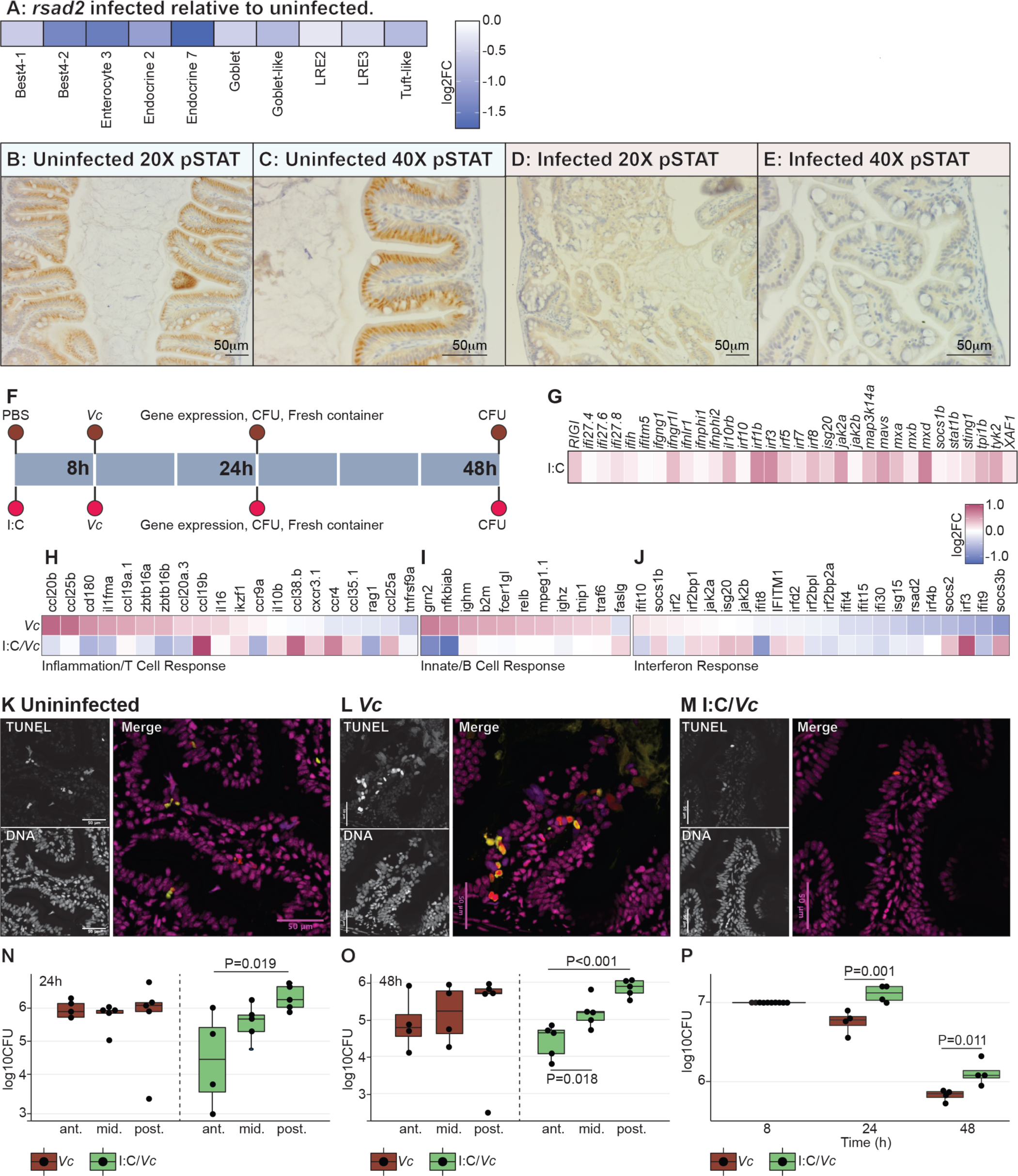
*Vc* Suppresses Host Interferon Signaling. **(A)** Heatmap for the expression levels of *rsad2* in mature intestinal epithelial cells. (B-E) Immunohistochemical imaging of phosphorylated STAT in the intestinal epithelium of uninfected (B-C) and infected (D-E) fish at the indicated magnifications. Scale bars are shown for each panel. (F) A timeline for quantitative NanoString-based gene expression analysis and water CFU levels of *Vc* in infected fish pre-treated without (brown circles), or with poly I:C (red circles). Each vertical while line represents an eight-hour time interval. Water changes were performed at 24h. (G-J) Heatmaps showing relative induction of indicated gene categories in fish treated with poly I:C, infected fish (*Vc*), and in infected fish pre-treated with poly I:C (I:C/*Vc*). (K-M). Visualization of TUNEL-positive cells in the intestinal epithelium of uninfected fish (K), fish challenged with *Vc* (L), and fish treated with poly I:C prior to infection (M). In the merged images, TUNEL+ cells are yellow, and nuclei are magenta. (N-O) Regional distribution of *Vc* in the intestines of fish challenged with *Vc* or pretreated with poly I:C prior to infection. (P) CFU levels of *Vc* in water sampled from fish treated with *Vc* or poly I:C prior to *Vc* for the indicated periods. For N-P, P values were calculated using an ANOVA test with Tukey’s corrections for multiple comparisons.

We consider possible interactions between *Vc* and host interferon intriguing, as interferon is the subject of natural selection in a human population where *Vc* is endemic^34^. Therefore, we examined the relationship between interferon activation, gut immunity, and disease severity in infected fish. In initial experiments, we used quantitative gene expression analysis to compare the intestinal immune response of infected adults (*Vc*) to an experimental group that we pre-treated with the interferon pathway agonist poly I:C prior to infection (I:C/*Vc*, Figure 6F). We confirmed that pre-treatment with poly I:C alone was sufficient to induce an interferon response in the intestines of wildtype adult fish (Figure 6G). In agreement with our single cell data, we found that infection with *Vc* alone drove an immune response characterized by induction of pro-inflammatory cytokines (e.g. *ccl25b*, *ccl19a.1*, *ccl19b* and *ccl20a.2*, Figure 6H), enhanced expression of genes associated with macrophage function (e.g. *grn2*, *fcer1gl*, *mpeg1.1* and *traf6*, Figure 6I), and supported B cell activation marked by production of the IgM heavy chain, *ighm*, and the IgA-related IgZ heavy chain, *igzh* (Figure 6J). Notably, we confirmed that infection with *Vc* greatly attenuated expression of numerous interferon response genes (Figure 6J). Thus, the data in Figure 6B-J confirm that *Vc* infection establishes a host intestinal environment that includes broad suppression of interferon-response genes.

From there, we quantified the impacts of pretreatment with poly I:C on the gut response to *Vc*. We found that poly I:C blocked *Vc*-dependent induction of many pro-inflammatory cytokines (e.g. *ccl20b*, *ccl25b* and *il1fma*, Figure 6H), and prevented *Vc*-mediated expression of critical signatures of B cell activation (e.g. *ighm*, *b2m*, *relb*, *ighz* Figure 6I). Instead, exposure to poly I:C enhanced expression of multiple IFN pathway components (e.g. *jak2a*, *jak2b*, *irf3* Figure 6J), and increased expression of a number of interferon-response genes (e.g. *isg20*, *ifitm1*, *rsad2*, *socs2*, *socs3b*, Figure 6J), indicating that poly I:C treatment tilted the gut *Vc* response towards an interferon signature.

To test if pre-treatment with poly I:C impacts *Vc*-mediated pathology, we then looked at cell death in intestines of adult fish that we pre-treated with poly I:C for five hours, followed by a sixteen-hour infection with *Vc*. In the absence of infection, we detected a limited number of TUNEL+ apoptotic cells in the gut (Figure 6K). Consistent with data from Figure 4, infection disrupted epithelial integrity, and increased the amount of dying cells, including material shed into the lumen (Figure 6L). In contrast, pre-treatment of fish with poly I:C restored epithelial integrity, and greatly diminished the appearance of TUNEL+ cells (Figure 6M), indicating that a poly I:C treatment protects the host epithelium for *Vc-*dependent damage.

We then used the infection paradigm outlined in Figure 6F to test if pre-exposure to poly I:C impacts intestinal colonization by *Vc*. Consistent with Figure 4A-C, we found that infection alone resulted in approximately equal loads of *Vc* in each intestinal segment at 24h and 48h (Figure 6N-O). In contrast, poly I:C-treated fish had a marked shift in the intestinal distribution of *Vc*. At both 24h and 48h time points, we observed a significant decline in the anterior *Vc* burden, with a marked increase in posterior accumulation of *Vc* (Figure 6N-O). A recent study showed that diminished anterior colonization by commensal *Aeromonas* leads to increased fecal shedding of the microbe^47^. Therefore, we asked if poly I:C exposure also impacted host shedding of *Vc*. We were particularly interested in effects on *Vc* shedding, as fish are candidate vectors for *Vc* transmission in the wild. For this assay, we challenged poly I:C-treated fish with *Vc* for 16h, transferred infected fish to a fresh container daily, and measured water contamination by *Vc* at 24h intervals (Figure 6F). At both 24 and 48h post infection, we found that pre-treatment with poly I:C significantly increased the shedding of *Vc* relative to untreated, infected controls (Figure 6J). Thus, our data indicate that *Vc* suppresses interferon signaling in an infected fish, and that induction of the interferon response diminishes host epithelial death, destabilizes anterior intestinal colonization by *Vc*, and significantly increases the release of live bacteria by infected animals.

## DISCUSSION

Physiological, genetic and developmental studies demonstrated extensive similarities between fish and mammalian intestines, highlighting the utility of zebrafish to characterize drivers of intestinal illness^25,48–52^. Due to *ex utero* development, genetic accessibility, and optical transparency, larvae provide a simple, manipulable system to track intestinal function in real time. However, the larval gut is small, has not completed development, and interacts with a limited repertoire of leukocytes^46^. In contrast, the large, structurally elaborate adult intestine associates with a complex leukocyte mix that includes critical immune-regulatory myeloid and lymphoid lineages. However, we lack a comprehensive picture of IEC composition and distribution in adult guts, and most studies of gut-associated leukocytes examined purified subsets that expressed fluorescent marker proteins. We consider a thorough, functional characterization of the adult gut essential for experimental manipulation of zebrafish as an intestinal disease model. To understand how the gut functions during homeostasis and in response to infection, we prepared single cell gene expression atlases of adult intestines that we raised under conventional conditions, or after infection with *Vibrio cholerae*. Our study identifies the adult gut as a sophisticated organization of specialist cells that collectively protect from damage, while maximizing nutrient acquisition.

Upon examination of gene expression data in conventional adults, we discovered that the gut contains tuft and Best4+ cells that were only recently identified in humans and developing zebrafish larvae^41,53,54^. The role of Best4+ cells in gut health is unclear. However, excess Best4 activity appears to support progression and metastasis of colorectal cancer^55^, suggesting a role for Best4+ cells in intestinal homeostasis. Mice lack Best4+ cells, preventing us from studying this cell type in a popular vertebrate model. However, with the extensive toolset available for genetic studies, we anticipate that zebrafish will emerge as a critical model to understand the impacts of Best4+ cells on health and disease. Likewise, our identification of IECs that express classical tuft cell markers establishes the adult as a powerful tool to explore the role of tuft cells in intestinal immunity, including in the context of gut-resident innate and adaptive immune cells. We also uncovered two distinct populations of cycling IECs: an undifferentiated cluster that expressed classical stem cell markers, alongside a second group enriched for Notch pathway components. We note similarities between the cycling cell populations of the zebrafish gut and the progenitor compartment of mammalian intestines formed by spatially restricted stem cells and more prevalent transit amplifying cells, and we provide supporting immunohistochemical evidence that the fish gut epithelium contains an abundance of proliferative cells at each fold base. However, we also caution that additional research is needed to clarify the exact identity of the cycling populations in the adult zebrafish gut.

We also discovered striking specializations within mature epithelial cell types, including enteroendocrine cells, enterocytes, and lysosome-rich enterocytes. Each lineage had transcriptionally distinct subsets that allow fine-tuned control of nutrient processing and interorgan communication. For example, we identified specialist enterocytes that are hallmarked by active interferon signaling, a critical regulator of IEC survival and proliferation^56,57^. Our work also expands prior transcriptional characterizations of intestinal blood cell types by simultaneously generating an unbiased expression atlas of all gut-associated leukocytes in uninfected and infected adults. We confirmed polarized macrophages^46^, intestinal T cells^42^, and innate lymphoid cells^42^, but also generated gene expression atlases for intestinal dendritic cells, B cells and granulocytes. Notably, we found that the adult intestine contains previously uncharacterized lymphoid aggregate-like structures with enriched populations of immune cells. Our work aligns with recent identification of mucosal-associated lymphoid tissue in the gill^58,59^ and suggests that fish leukocyte defenses are elaborate in terms of organization and contribution from participant cells.

Alongside a transcriptional profile of conventional guts, we resolved the host response to *Vc* to the level of individual intestinal cell types. *Vc* is a deadly, non-invasive pathogen that impacts human populations globally. Fish are established models of *Vc* pathogenesis^24,60,61^, but we lack detailed understanding of the impacts of a natural, primary encounter with *Vc* in patients or biomedically relevant models. Our work supports patient data that *Vc* causes a mild, inflammatory response that includes T cell engagement, but also clarifies the impact of *Vc* on host IECs, as well as gut-associated leukocytes. We discovered a coordinated response of IECs and their associated lymphocytes to infection that resulted in attenuated interferon signaling, and a moderate immune response that included initiation of T cell-based defenses.

We consider interferon inhibition a striking feature of the fish response to *Vc*. Type I interferons act directly on the epithelium to arrest proliferation and prevent apoptosis, thereby supporting cellular maturation and maintaining epithelial barrier integrity^62,63^. At the same time, interferon acts on gut-associated leukocytes, including Treg, Th1 and Th17 cells to suppress inflammation and contribute to the repair of damaged epithelia^64–66^. In this regard, it is interesting that interferon appears to be the subject of natural selection in a human population where *Vc* is endemic, raising the possibility that modified interferon signaling impacts the progression of a *Vc* infection. Consistent with this hypothesis, we found that exposure to an interferon pathway agonist impacted colonization by *Vc*, protected IECs from *Vc*-dependent cell death, and enhanced the diarrheal shedding of live pathogen by infected fish. Although, additional work is necessary to understand how interferon impacts gut defenses, we believe this study provides a rational cellular roadmap to track the involvement of each gut cell type in host intestinal immune response to *Vc*.

## MATERIALS AND METHODS

### Animals

All zebrafish experiments were performed at the University of Alberta using protocols approved by the University’s Animal Care & Use Committees, Biosciences and Health Sciences (#3032), operating under the guidelines of the Canadian Council of Animal Care. Wild type TL zebrafish were reared at 29°C in tank water (Instant Ocean Sea Salt dissolved in reverse osmosis water for a conductivity of 1000µS and pH buffered to pH 7.0 with sodium bicarbonate) under a 14h/10h light/dark cycle using standard zebrafish husbandry protocols. Zebrafish were fasted for 20h prior to infections.

### Generation of transgenic zebrafish

Zebrafish with the promoter region of the pan-phagocyte marker, mpeg1.1^67^, or T-cell marker, lck (Jeffrey Yoder, unpublished), driving EGFP expression were generated using the Tol2kit^68^. Entry plasmids were purchased from Addgene (p5E-mpeg1.1 plasmid #75023 and p5’Entry-lck-5.5kb plasmid #58890) then recombined into pDestTol2CG2 along with pME-EGFP and p3E-polyA using gateway cloning. The final constructs were confirmed via restriction digest and co-injected with transposase RNA into one-cell stage, wild type TL embryos. Larvae were screen for *cmlc2:EGFP* and reporter co-expression. A stable line was established by outcrossing founders to wild type AB zebrafish and screening for GFP expression.

### Infection with *Vibrio cholerae* by immersion

The V52 *V. cholerae* strain was streaked from a glycerol stock onto LB agar supplemented with streptomycin (100µg/mL final) and grown overnight at 37°C. A single colony was grown overnight with aeration at 37°C in LB broth supplemented with streptomycin (100µg/mL final). Bacteria were washed with 1xPBS (pH 7.4) then resuspended to an OD600 of 1 (∼10^8^CFU/mL) in 1xPBS. Five fish were incubated in a 400-mL beaker containing 1mL of the OD600 = 1 culture or 1mL of 1xPBS (‘uninfected’) in 200mL sterile tank water (filtered through a 0.22µm filter) at 29°C and 14h/10h light/dark cycle as described previously^21^. To enumerate *V. cholerae* CFU in the intestine and tank water over time, fish were infected with GFP-expressing strains and fluorescence used to count *V. cholerae* colonies. Fish were transferred to fresh, sterile tank water and fed daily until the end of each experiment.

### Treatment of zebrafish with Poly I:C by immersion

Five zebrafish housed in 400-mL beakers containing 200mL sterile tank water were immersed in high molecular weight polyinosine-polycytidylic acid (HMW poly I:C, InvivoGen Cat.# tlrl-pic-5) (50µg/mL final concentration in 200mL tank water) or 1xPBS (‘untreated’) for 5h. Five hours after treatment, fish were infected with *V. cholerae* or treated with 1xPBS, as indicated above, for 16h.

### Generating single-cell suspensions for single cell RNA-sequencing

Five fish were infected or treated with PBS for 16h as described above. Zebrafish intestines were dissected, surrounding organs removed, then minced into smaller pieces with Vannas scissors. Minced intestinal tissue was washed in 10mL PBS (5 minutes, 300rcf, 4°C). The tissue was digested in a dissociation cocktail containing fresh collagenase A (1mg/mL), 40µg/mL Proteinase K, and Tryple (Gibco 12605-010, diluted 1:1000 [final]) in PBS for 30 min in a 37°C water bath. Tissue was mechanically disrupted by gentle pipetting 40X every 10 minutes to aid in digestion. A 10% stock of non-acetylated bovine serum albumin (BSA) was added at a final concentration of 1% in PBS to stop the dissociation after 30 minutes. Cell suspensions were diluted in 5mL ice-cold PBS then filtered through a 40µm filter fitted on a 50mL conical. Cells were pelleted (15min, 300rcf, 4°C) then the pellet was resuspended in 450µL PBS + 0.04% BSA and live cells collected using OptiPrep™ Density Gradient Medium (Sigma, D1556-250ML). OptiPrep™ Application Sheet C13 was followed to remove dead cells. Briefly, a 40% (w/v) iodixanol working solution was prepared by diluting 2 volumes of OptiPrep™ in 1 volume of PBS + 0.04% BSA. The working solution was used to prepare a 22% (w/v) iodixanol solution in PBS + 0.04% BSA. One volume of working solution was carefully mixed with 0.45 volume of cell suspension by gentle inversion. The cell suspension mixture was transferred to a 15mL conical tube and topped up to 6mL with working solution. The working solution was then overlaid with 3mL 22% (w/v) iodixanol diluted PBS + 0.04% BSA. Finally, the 22% iodixanol solution was overlaid with 0.5mL PBS + 0.04% BSA. Live cells were separated by centrifuging at 800g for 20 minutes at 20°C. Viable cells were collected from the top interface (top 500µL). Cells were washed in 1mL PBS + 0.04% BSA (10min, 300rcf, 4°C), supernatant removed, then resuspended in 50µL PBS + 0.04% BSA. Cell suspensions were filtered through a 40µm filter fitted on a 1.5mL tube. The filter was washed with 20µL PBS + 0.04% BSA. Viability and cell counts were determined by staining dead cells with Trypan blue and counting with a hemacytometer. Cell suspensions were then diluted to a concentration of 1200 cell/µL and immediately run through the 10X Genomics Chromium Controller with Chromium Single Cell 3’ Library & Gel Bead Kit v3.1. Library QC and sequencing was performed by Novogene using the Illumina HiSeq platform.

### Processing and analysis of single-cell RNA-seq data

Cell Ranger v3.0 (10X Genomics) was used to demultiplex raw base call files from Illumina sequencing and to align reads to the Zebrafish reference genome (Ensembl GRCz11.96). Cell Ranger output matrices were analyzed using the Seurat R package version 4.1^69^ in RStudio. Cells possessing fewer than 200 unique molecular identifiers (UMIs), greater than 2500 UMIs, or greater than 25% mitochondrial reads were removed to reduce the number of low-quality cells and doublets. Seurat was then used to normalize expression values and perform cell clustering on individual or integrated datasets at a resolution of 0.8 with 50 principal components (PCs), where optimal PCs were determined using JackStraw scores^70^ and elbow plots. After using the “FindMarkers” function in Seurat to identify marker genes for each cluster, clusters were annotated according to known cell type markers in zebrafish, or orthologous markers in mammals.

### RNA extraction for Nanostring

RNA was extracted from zebrafish intestinal tissue using TRIzol™ Reagent (Invitrogen Cat# 15596026) as follows. Three zebrafish intestines were dissected, surrounding tissue removed, then quickly homogenized in 250µL TRIzol™. The intestinal homogenates were topped up to 1mL with TRIzol™. Homogenized tissue was stored at −70°C prior to RNA extraction. Four hundred microliters of chloroform were added to destabilize the aqueous phase, gently vortexed for 15 seconds then incubated at room temperature for 3 minutes. Chloroform extraction was repeated with the upper aqueous phase at a 1:1 ratio. Sodium acetate (3M, pH 5.2) was added at one tenth the volume of the collected aqueous phase (50µL sodium acetate). Ninety-five percent ethanol was added at two times the volume (1mL) then incubated overnight at 4°C to precipitate salts and nucleic acids. Nucleic acids were pelleted by centrifuging at 12,000rcf, 10min 4°C. Ethanol was carefully removed. RNA pellet was washed twice with 0.5mL 75% ethanol (7500rcf, 10min, 4°C). RNA pellet was allowed to dry 4 minutes then resuspended in 50µL nuclease-free water. RNA concentrations were determined by Nanodrop then diluted to 10µL at 100ng/µL for Nanostring hybridization using the nCounter® Elements™ platform.

### Immunofluorescence and fluorescence microscopy

Zebrafish intestines were fixed in 4% PFA (EMS Cat #15710, 16% paraformaldehyde diluted in 1xPBS) overnight at 4°C. Intestines were washed twice in 1xPBS. Posterior intestinal segments were cut from the rest of the intestine then cryoprotected in 15% (w/v) sucrose/PBS at room temperature until sunk followed by sinking in 30% (w/v) sucrose/PBS (overnight at 4°C). Posterior intestines were mounted in Optimal Cutting Temperature Embedding Medium (Fisher Scientific Cat#23-730-571), then frozen on dry ice. Five-micron cryosections were collected on Superfrost Plus (Fisherbrand Cat# 22-037-246) slides. After allowing sections to adhere onto slides, slides were immersed in PBS for 20 minutes at room temperature. Tissue was blocked for 1h at room temperature in 3% (w/v) BSA dissolved in PBST (1xPBS + 0.2% (v/v) Tween-20). Primary antibodies were diluted in blocking buffer and layered onto sections. Sections were incubated in primary antibody solution overnight in a humid chamber at 4°C. Excess primary antibody was washed by immersing slides in fresh PBST for 1.5h. Sections were incubated in secondary antibody solution (prepared in blocking buffer) for 1h at room temperature in a humid chamber, protected from light. Secondary solution was removed then sections incubated in Hoechst (Molecular Probes Cat# H-3569) diluted 1:2000 in PBST for 10 minutes protected from light. Slides were washed in PBST by brief immersion followed by a 30-minute incubation in fresh PBST protected from light. Slides were mounted in Fluoromount™ Aqueous Mounting Medium (Sigma F4680-25ML).

### Hematoxylin & Eosin staining

Zebrafish intestines were fixed in 4% neutral buffered formalin at room temperature. Posterior intestinal segments were processed for paraffin embedding and 5µm sections collected on slides. Tissue was deparaffinized then stained with Hematoxylin Gill III (Leica Ca.# 3801542) for two minutes followed by an eosin counterstain (Leica Cat# 3801602). Sections were dehydrated, cleared in toluene, then mounted in DPX (EMS Cat#13512).

### Immunohistochemistry

Zebrafish intestines were fixed at 4°C in BT fixative: 4% PFA, 0.15mM CaCl2, 4% Sucrose in 0.1 M phosphate buffer (pH 7.4)^26^. Posterior intestinal segments were processed for paraffin embedding and 5µm sections collected on Superfrost Plus slides. Tissue was deparaffinized, rehydrated, then boiled in 10mM sodium citrate pH 6, 0.05% Tween-20 for 20 min at 98°C to unmask antigen. After cooling sections to room temperature for 30min, sections were incubated in 3% hydrogen peroxide for 10 minutes, washed twice in dH_2_O, and once in PBST (PBS + 0.5% Triton-X 100). Sections were then blocked for 1h at room temperature in 10% (v/v) normal goat serum (NGS)/PBST followed by overnight incubation in primary antibody solution (prepared in blocking buffer) in a humid chamber at 4°C. Slides were washed three times in PBST and tissue incubated in SignalStain® Boost Detection Reagent (HRP, Rabbit or Mouse) for 30 min at room temperature. Colorimetric detection was done by incubating in SignalStain® DAB Chromogen solution (Cell Signaling Technology #8059) for 5 minutes. Sections were counterstained in ¼-strength Hematoxylin Gill III for 30 seconds dehydrated in ethanol, cleared with toluene and mounted in DPX.

### HCR™ RNA fluorescence *in situ* hybridization

Whole zebrafish intestines were fixed in 4% PFA in PBS overnight at 4°C, then washed in PBS. Intestines were then positioned in OCT (Fisher Scientific 4585) in base molds (Fisher Scientific 22-363-552) and frozen on dry ice. After tissue sectioning (5 µm sections), slides were stored at −70°C until processing. Tissue was then prepared according to established protocols (Molecular Instruments). Briefly, tissue was immersed in ice-cold 4% PFA for 15 min at 4°C. Tissue was then dehydrated with a graded ethanol wash series (50%; 70%; 100%; 100%), followed by two PBS washes. Slides were then pre-hybridized with probe hybridization buffer (Molecular Instruments) at 37°C for 15 minutes inside humid chamber. Next, tissue was incubated in a humid chamber overnight at 37°C with probes in hybridization buffer and covered with parafilm to prevent liquid loss. Probes were then removed, and slides were washed with probe wash buffer (Molecular Instruments), followed by a series of 15-minute washes at 37°C with probe wash buffer/ 5X SSCT (75/25; 50/50; 25/50; 0/100). After an additional wash with 5X SSCT (0.1% Tween-20) at room temperature, slides were incubated with amplification buffer (Molecular Instruments) for 30 minutes at room temperature. At the same time, hairpin amplifiers were heated to 95°C for 90 seconds and snap-cooled to room temperature for 30 minutes. Hairpins were mixed in amplification buffer, and slides were incubated with hairpin solution (covered with parafilm) overnight in a humid chamber at room temperature. The next day, slides were washed in 5X SSCT and counter-stained with Hoechst 33258 (Molecular Probes Cat# H-3569) diluted 1:2000 in 5X SCCT before mounting in Fluoromount™ Aqueous Mounting Medium (Sigma F4680-25ML).

### Intestinal leukocyte isolation

Freshly dissected zebrafish intestines were gently squeezed between the frosted ends of two glass slides while immersed in 1xPBS pH 7.4, 25mM Hepes, 1mM EDTA, 2% heat inactivated new born calf serum (Gibco 26010-066). The tissue/cell suspension was filtered through a 70µm filter to remove large tissue chunks. Single cells were pelleted (300rcf, 10min, 4°C), supernatant was decanted, and cells resuspended in remaining liquid. For flow cytometry, nuclei were stained with DRAQ5 (Invitrogen 65-0880-96). Prior to analysis by flow cytometry or cytospin, cells were filtered through a 40µm filter to remove large cells. Unfixed, nucleated, GFP-positive cells were analyzed on an Attune NxT (Life Technologies) or sorted on a Sony MA900. Cell suspension and sorted cells were cytospun (ThermoShandon Cytospin4) onto a Superfrost Plus slide for 3min at 10g^71^. Cells were allowed to air dry then fixed with methanol. Cells were stained with May-Grünwald solution (Sigma 63590) for 10 minutes and counterstained with Giemsa (Sigma 48900) for 45 minutes then mounted.

### Microscopy

Histological samples were imaged on a ZEISS AXIO A1 compound light microscope with a SeBaCam 5.1MP camera. Confocal images were captured on an Olympus IX-81 microscope fitted with a CSU-X1 spinning disk confocal scan-head, Hamamatsu EMCCD (C9100-13) camera and operated with Perkin Elmer’s Volocity. Confocal Z-stack image processing was done in Fiji. May-Grünwald-Giemsa-stained leukocytes were imaged on a Zeiss AxioObserver.Z1 with an Axiocam HRm CCD camera.

### Transmission electron microscopy of *V. cholerae* in adult zebrafish intestines

Zebrafish were infected with *V. cholerae* strain V52 for 16h by immersion as described above. Posterior intestines were fixed, processed, and imaged as described in^41^.

### Additional resources

Processed single-cell RNA-seq data for uninfected adult fish is available for visualization and analysis on the Broad Single Cell Portal: https://singlecell.broadinstitute.org/single_cell/study/SCP2141/adult-zebrafish-intestine#studyvisualize. Single-cell RNA-seq raw data files for uninfected and infected fish have been deposited to the NCBI GEO database and are publicly available (GSE230044: https://www.ncbi.nlm.nih.gov/geo/query/acc.cgi?acc=GSE230044).

## SUPPLEMENTARY FIGURES

**Supplementary Figure 1:**
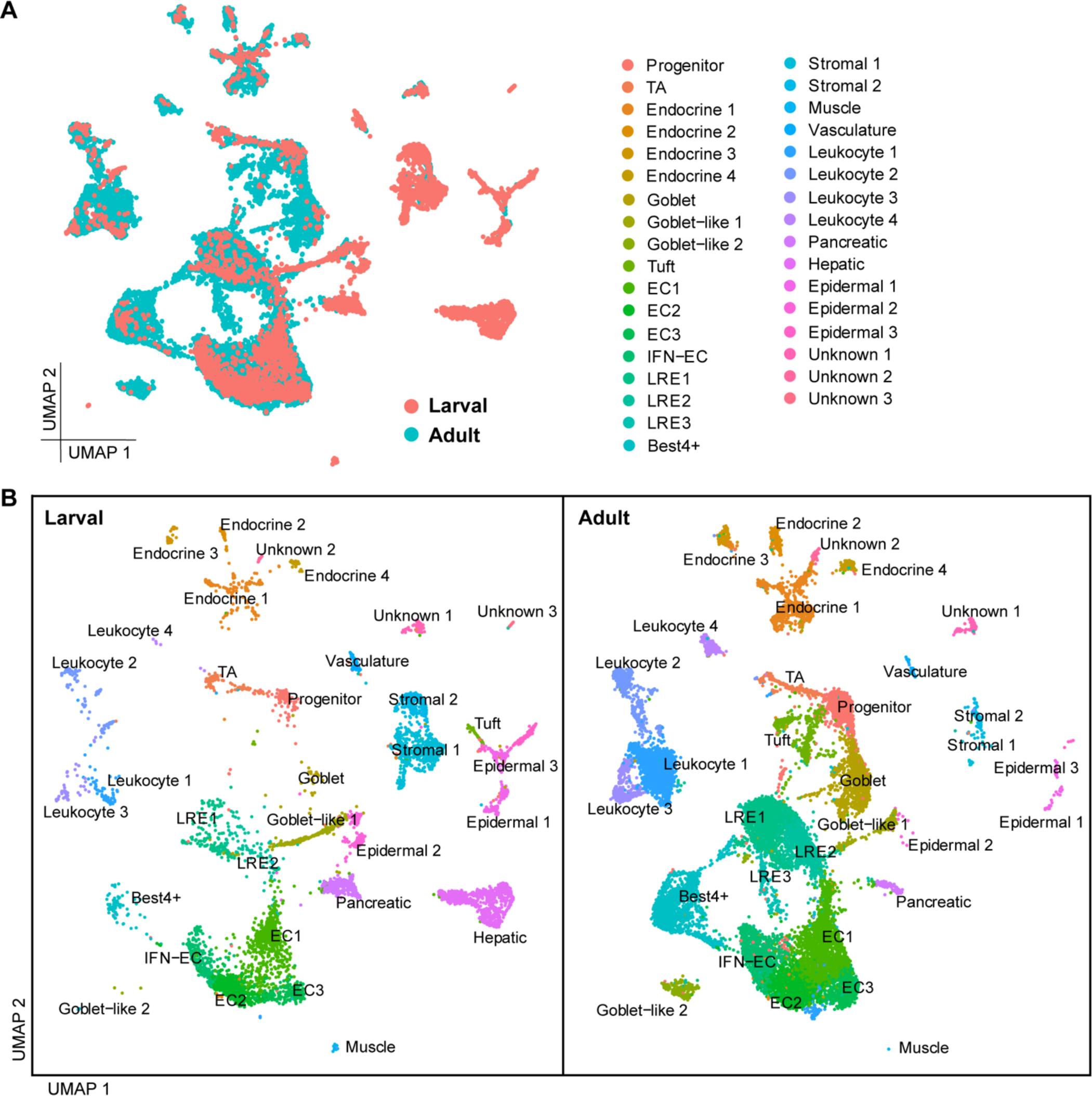
Comparison of transcriptional states in larval and adult intestines. (A) 2D t-SNE projections of integrated single cell gene expression data from larvae (orange) and adults (cyan). (B) 2D t-SNE projections of individual larval and adult data sets with cell types color-coded for each set.

**Supplementary Figure 2:**
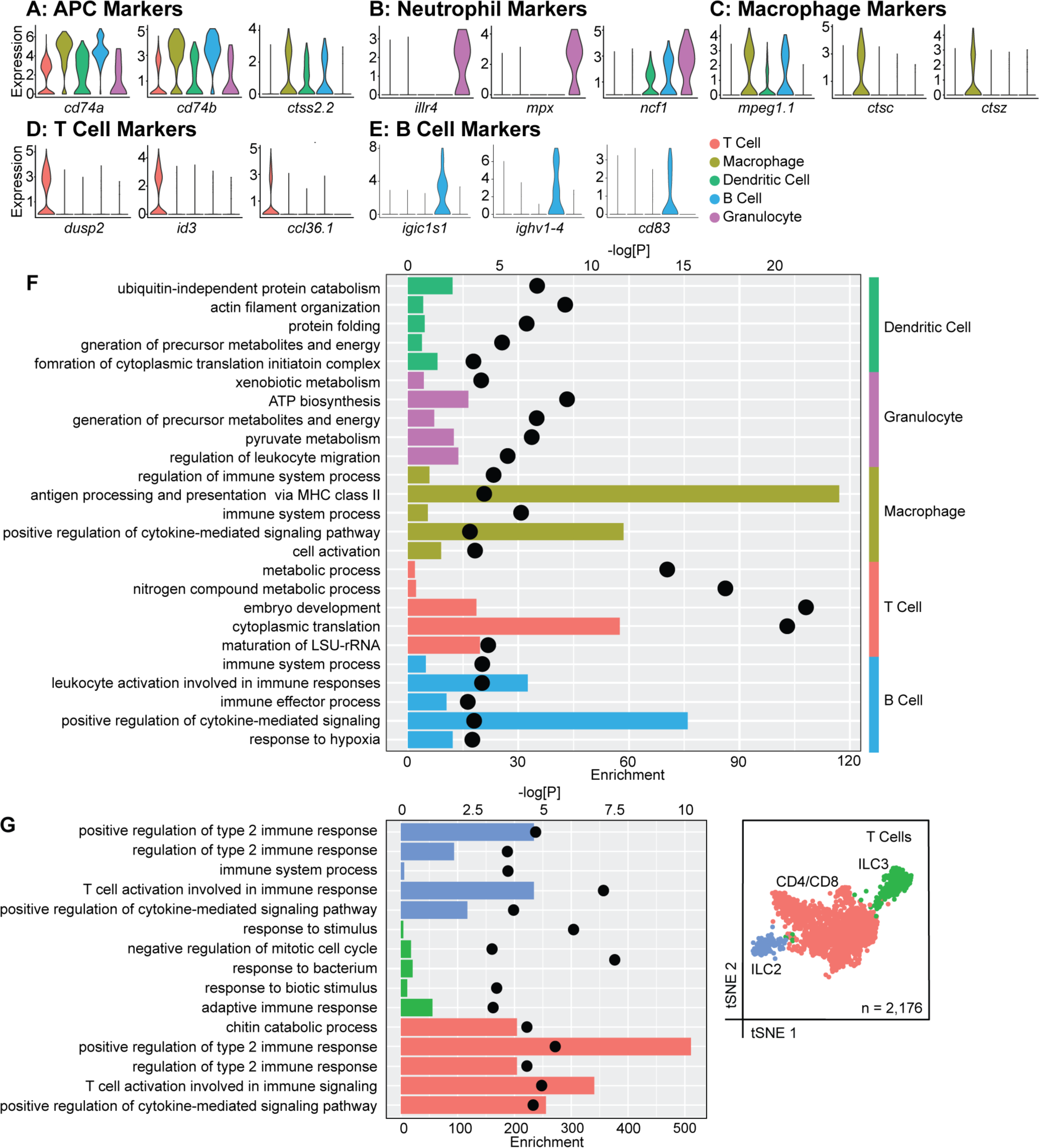
Transcription Characterization of Gut-Associated Leukocytes. (A-E) Relative expression of the indicated lineage-specific markers in gut-associated leukocytes. (F) The top five enriched non-redundant Gene Ontology terms per cell type. Enrichment score is represented by bar length and the negative log p value is shown with black circles. (G) The top five enriched non-redundant Gene Ontology terms associated with intestinal T cells and type 2 and 3 ILCs. Enrichment score is represented by bar length and the negative log p value is shown with black circles.

**Supplementary Figure 3:**
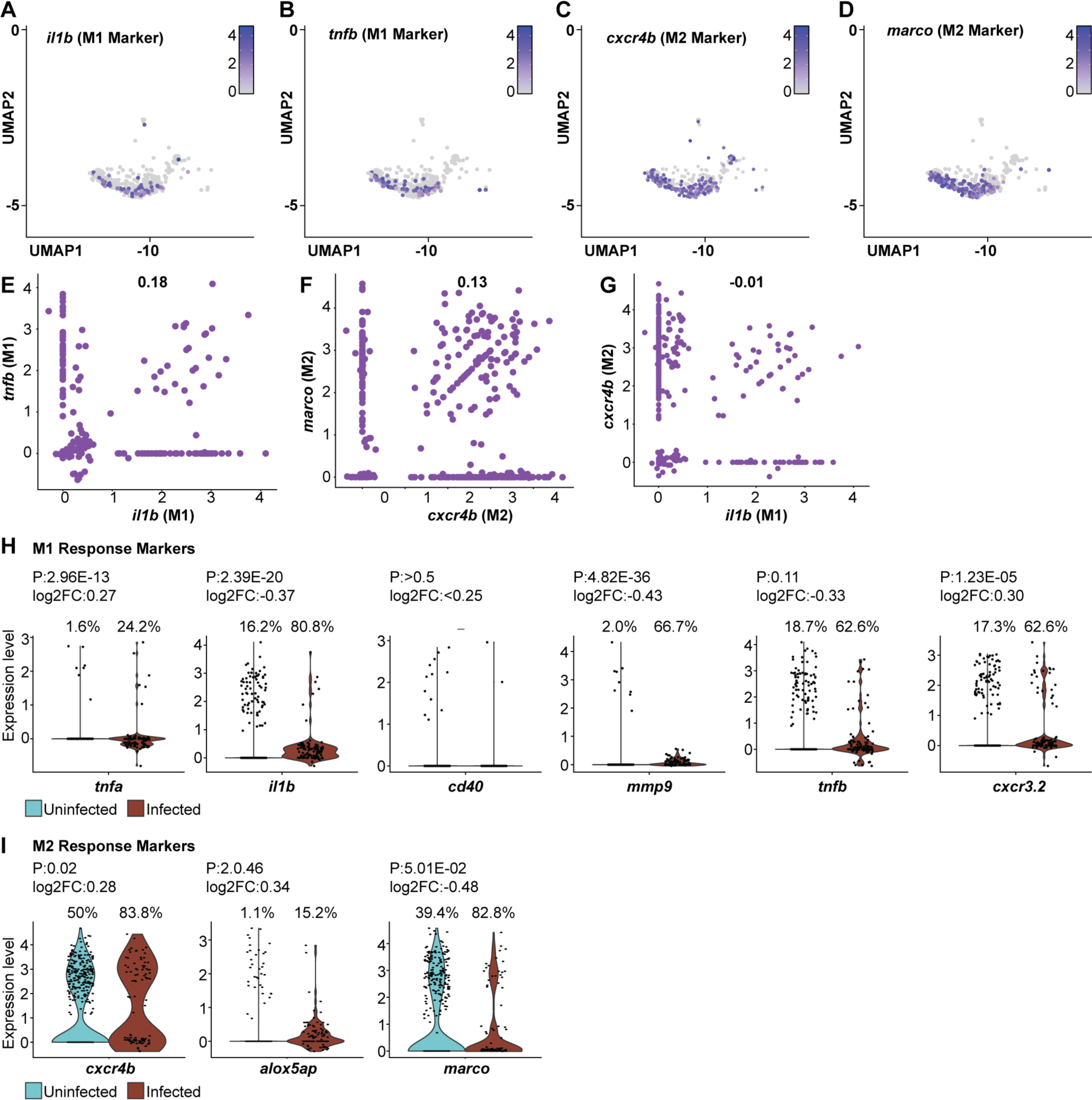
Transcription Characterization of Gut-Associated Macrophages. (A-D) Relative expression of the indicated M1 and M2 markers in gut-associated macrophages. (E-G) Scatter plots examining the relative co-expression of M1 markers (E) M2 markers (F) and M1 and M2 markers (G) in conventional Macrophages. r value indicates Pearson correlation for all three plots. (H-I) Relative expression of M1 response markers and M2 responses markers in macrophages of uninfected (teal) and infected (brown) fish. For each transcript, P values and log2 fold-change in infected fish is shown.

## ACKNOWLEDGMENTS

Imaging experiments were performed at the University of Alberta Faculty of Medicine & Dentistry (FoMD) Cell Imaging Core, RRID:SCR_019200, which receives financial support from FoMD, the Department of Medical Microbiology and Immunology, the University Hospital Foundation, and Canada Foundation for Innovation (CFI) awards to contributing investigators. TEM experiments were performed at the Department of Oncology Cell Imaging Facility. Flow Cytometry experiments were performed at the FoMD Flow Cytometry Facility, RRID:SCR_019195, which receives financial support from FoMD, the Li Ka Shing Institute of Virology, and Canada Foundation for Innovation (CFI) awards to contributing investigators. Single-cell libraries were prepared at the FoMD High Content Analysis Core, RRID:SCR_019182, which receives financial support from FoMD, the Li Ka Shing Institute of Virology, and Canada Foundation for Innovation (CFI) awards to contributing investigators. Immunohistochemistry experiments were performed at the Faculty of Science Department of Biological Sciences Advanced Microscopy Facility. We would also like to thank Science Animal Support Services and the Alberta Health Sciences Animal Laboratory Services for their excellent care of the zebrafish aquatics facilities. This work was supported by grants from the Canadian Institutes of Health Research (grant no. MOP77746) and the Weston Family Microbiome Initiative to EF. RJW has funding support through the National Science and Engineering Research Council Graduate Scholarships, and Alberta Innovates Graduate Student Scholarships.

